# Programmable macromolecule delivery via engineered trogocytosis

**DOI:** 10.1101/2025.03.12.642522

**Authors:** Xinyi Chen, Yinglin Situ, Yuexuan Yang, Maylin Lum Fu, Luna Lyu, Lei Stanley Qi

**Affiliations:** Department of Bioengineering, Stanford University, Stanford, CA 94305, USA; Institute for Computational and Mathematical Engineering, Stanford University, Stanford, CA 94305, USA; Sarafan ChEM-H, Stanford University, Stanford, CA 94305, USA; Chan Zuckerberg Biohub – San Francisco, San Francisco, CA 94108, USA

## Abstract

Trogocytosis, the transfer of plasma membrane fragments during cell-cell contact, offers potential for macromolecular delivery but is limited by uncertain fate of trogocytosed molecules, constraints to membrane cargo, and unclear generalizability. Here, we demonstrate that donor cells engineered with designed receptors specific to intrinsic ligands can transfer proteins to recipient cells through direct contact. We identified key principles for enhancing contact-mediated transfer and subsequent functionalization of transferred macromolecules, including receptor design, pH-responsive membrane fusion, inducible cargo localization, release, and subcellular translocation. Exploiting these findings, we developed TRANSFER, a versatile delivery system that integrates logic gate-based control to sense multiple ligand inputs and deliver diverse functional cargos for genome editing and targeted cell ablation across cell types. The study establishes trogocytosis as a novel, programmable framework for cell-based macromolecular delivery.

## Introduction

Cells have evolved intricate mechanisms for direct material transfer as an approach of cell-cell communication. While intercellular transport often relies on secreted compartments like extracellular vesicles, which have inspired methods of functional cargo delivery (*1*), trogocytosis represents a distinct, contact-dependent process that shows promise for molecular delivery.

During trogocytosis, one cell extracts plasma membrane fragments and associated molecules from another and incorporates them onto its own surface (*2*). Since the first report in the 1970s of intercellular transfer of major histocompatibility complex (MHC) between immune cells (*3*), trogocytosed proteins have been shown to retain function upon acquisition across disparate contexts, primarily in immune cells (*4*). For instance, chimeric antigen receptor (CAR) T cells can acquire antigens from cancer cells, reducing antigen density on tumor cells and leading to T cell exhaustion or fratricide (*5*). Similarly, natural killer cells that acquire CD9 (*6*) or PD-1 (*7*) via trogocytosis can exhibit diminished anti-tumor responses.

Unlike many intercellular transfer mechanisms, trogocytosis can exhibit high specificity, driven by ligand-receptor interactions. For example, T cell receptor (TCR) recognition of peptide-MHC molecules can initiate membrane protein transfer to target cells via trogocytosis (*8*). Despite these observations, the potential of trogocytosis as a delivery platform remains largely unexplored. This is due to a limited understanding of the fate and functional consequences of trogocytosed proteins in recipient cells, particularly non-immune cells (*9*); the requirement of cargo to be membrane proteins; and the lack of receptor designs capable of facilitating efficient intercellular transfer.

Overcoming these limitations could establish trogocytosis as a novel and generalizable mechanism for programmable, cell-specific molecular delivery using live cells. Unlike existing delivery methods that rely on synthetic materials such as lipid nanoparticles, viruses, virus-like particles, or extracellular vesicles (*1, 10*–*14*), engineered cells offer distinct advantages. They can continuously produce desired biomolecules, dynamically respond to external cues, and actively engage cellular pathways for targeted delivery to specific cells. Additionally, cell engineering allows for the incorporation of temporal control and logic-based integration of multiple ligand inputs, enhancing delivery precision and specificity.

Here we investigate the potential of trogocytosis as a programmable, versatile, and specific molecular delivery mechanism applicable to diverse cell types. We explore the molecular designs of trogocytosis receptors for selective recognition of intrinsic ligands on target cells and elucidate principles for functionalizing trogocytosed molecules to enable broad functions in recipient cells.

## Results

### Engineered receptors mediate highly specific cell-to-cell molecular transfer via trogocytosis

We started by co-culturing human Jurkat T lymphocytes stably expressing mCherry-tagged anti-CD19 CAR with K562 leukemia cells expressing GFP-tagged CD19 (**Fig. 1a**). To distinguish donor and recipient cells, recipient cells were engineered to stably express nuclear-localized BFP that cannot undergo intercellular exchange. After 24 hours of co-culture, we observed significant increase in mCherry in recipient cells, which correlated with CD19 expression levels (**Fig. 1b**).

**Figure 1:**
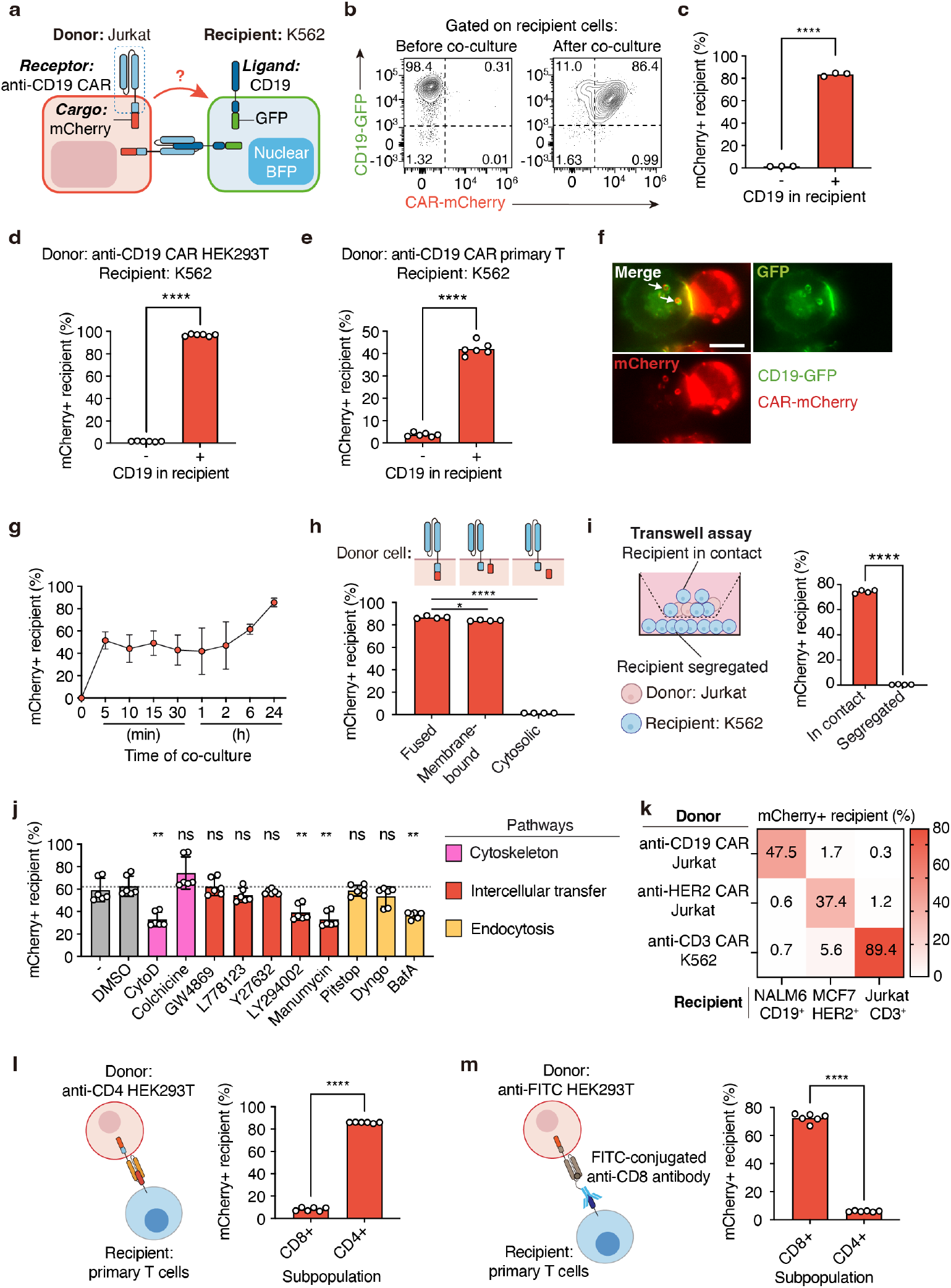
Engineered receptors mediate ligand-specific cell contact-based molecular transfer via trogocytosis. (**a**) Schematic illustrating molecule transfer via trogocytosis between donor Jurkat cells stably expressing an engineered receptor (anti-CD19 CAR-mCherry) and recipient K562 cells stably expressing the cognate ligand (CD19-GFP). mCherry and GFP were fused to the C-terminus of the respective transmembrane proteins, and nuclear-localized BFP was stably expressed only in recipient cells. (**b**) Flow cytometry before and after the 24-hour 1:1 co-culture of donor and recipient cells. Live, singlets, and BFP^+^ recipient cells were gated. Numbers indicate subpopulation frequency (%) in each quadrant. (**c**) Percentage of trogocytosed mCherry^+^ recipient cells among BFP^+^ recipient cells after 24-hour 1:1 co-culture of anti-CD19 CAR-mCherry Jurkat and CD19-GFP K562 cells, measured by flow cytometry. n = 3 biological replicates with mean. (**d**) Percentage of mCherry^+^ recipient cells upon co-culture between anti-CD19 CAR-mCherry HEK293T donor cells and K562 recipient cells at 5:1 for 24 hours as measured by flow cytometry. n = 6 biological replicates from 2 independent experiments. (**e**) anti-CD19 CAR-mCherry primary T cells were used as donor cells in a 1:1 donor-to-recipient cell ratio. Co-culture duration was limited to 6 hours to minimize cytotoxicity. n = 6 biological replicates from 2 human donors. Recipient cells were recognized as BFP^+^ populations. (**f**) Microscopy images of anti-CD19 CAR-mCherry Jurkat and CD19-GFP K562 cells during 1:1 co-culture at 9 minutes. Channels for mCherry, GFP, and merged images are shown. mCherry in donor cells was overexposed to highlight the weak signal in recipient cells. Scale bar, 10 μm. (**g**) Time-course flow cytometry of mCherry transfer over 24-hour co-culture. mean ± SD of n = 6 biological replicates across 2 independent experiments. (**h**) Top, schematic of mCherry localization in donor Jurkat cells. mCherry is membrane-tethered via a Myristoylation tag in group 2. Bottom, percentage of mCherry^+^ recipient cells after 24-hour 1:1 co-culture with different donor cells, measured by flow cytometry. n = 4 biological replicates. (**i**) Left, schematic of a transwell co-culture setup with CD19^+^ recipient cells seeded either in the same chamber (recipient in contact) and the opposite chamber (recipient segregated) as donor cells. Right, percentage of mCherry^+^ recipient cells after 24-hour 1:1 co-culture, measured by flow cytometry. n = 4 biological replicates. (**j**) Inhibitor effects on mCherry trogocytosis during 2-hour 1:1 co-culture of anti-CD19 CAR-mCherry Jurkat and CD19-GFP K562 cells. Mean ± SD of n = 6 biological replicates across 2 independent experiments. (**k**) Ligand-specific mCherry trogocytosis in a panel of donor and recipient cell types. Donor cells stably expressing anti-CD19 CAR, anti-HER2 CAR, or anti-CD3 CAR fused to mCherry were co-cultured with different recipient cells. Percentages of mCherry^+^ recipient cells (identified with nuclear BFP) after 24-hour co-culture are shown. Donor-to-recipient ratio 1:5 for anti-CD19 and anti-CD3 donors and 5:1 for anti-HER2 donors, determined as described in **Methods**. Mean of n = 6 biological replicates across 2 independent experiments. (**l, m**) Donor HEK293T cells expressing CAR-mCherry targeting CD4 or CD8 are co-cultured with primary T cells for 2 hours. Primary T cells are identified by antibody staining. Percentages of mCherry^+^ cells in CD8^+^ and CD4^+^ subsets are quantified. Donor-to-recipient ratios: 1:5 for anti-CD4 donors, 1:1 for anti-CD8 donors. n = 6 biological replicates from 2 human donors. Each dot represents one replicate culture well. Statistical tests: Welch’s two-sided t test for comparing 2 conditions. For inhibitor assay in **j**, Brown-Forsythe and Welch ANOVA test was used, with multiple comparisons to DMSO group. p<0.0001, ****; 0.0001≤p<0.001, ***; 0.001≤p<0.01, **; 0.01≤p<0.05, *; p≥0.05, non-significant (ns). Error bars indicate mean ± SD.

CAR-mCherry transfer was CD19-specific (**Fig. 1c**) and observed across other donor cell types, including HEK293T and primary T cells (**Fig. 1d, e**). Time-lapse microscopy showed CAR-mCherry signals appearing in recipient cells within 6 minutes of contact, colocalizing with CD19-GFP (**Fig. 1f**). Flow cytometry revealed that over 40% of recipient cells became mCherry^+^ within 5 minutes of co-culture (**Fig. 1g**). This rapid time scale is consistent with direct protein transfer, rather than DNA or mRNA transfer (*15*). Donor cells expressing membrane-bound mCherry not fused to CAR also transferred mCherry, while cytosolic mCherry showed negligible transfer, highlighting the preferential transfer of membrane-bound molecules during trogocytosis (*8*) (**Fig. 1h**).

To confirm that molecular transfer was not mediated by secreted vesicles, we conducted a transwell assay with 3 μm pores, which blocked direct cell-cell contact but allowed extracellular vesicles (30-1000 nm) to pass through (*16*). No mCherry transfer was detected in the segregated recipient cells, confirming that molecular transfer is cell-contact dependent (**Fig. 1i**).

Receptor trogocytosis was significantly inhibited by Cytochalasin D (CytoD), an actin polymerization blocker (*17*), but not by the microtubule inhibitor Colchicine (*18*) (**Fig. 1j**). This observation aligns with reports that TCR-mediated trogocytosis depends on actin polymerization (*7, 17, 19*). Inhibitors targeting Ras farnesylation (Manumycin-A) (*20*), phosphoinositide 3-kinase (PI3K) (LY294002) (*21*), and vacuolar H^+^ ATPase (Bafilomycin A1, BafA) (*22*) also significantly impaired transfer at non-cytotoxic concentrations. In contrast, inhibitors of exosome biogenesis (GW4869) (*23*), tunneling nanotube formation (L778123) (*24*), microvesicle biogenesis (Y27632) (*23*), clathrin-mediated endocytosis (Pitstop1), and dynamin-mediated endocytosis (Dyngo4A) (*25*) showed no effect.

To demonstrate the versatility of engineered trogocytosis across cell types and its specificity for endogenous ligands, we tested multiple recipient cell types, including NALM6 (B cell precursor leukemia), MCF7 (breast cancer), and Jurkat cells. RNA-sequencing datasets identified CD19, HER2, and CD3 as their respective surface markers (*26*). Co-culturing donor cells expressing CARs specific to these ligands with recipient cells in all combinations revealed highly specific transfer dependent on the cognate ligand (**Fig. 1k**).

We further evaluated the specificity of cell-to-cell transfer in primary T cells using various ligand-receptor pairs. Using HEK293T donor cell with mCherry as the cargo, we observed that donor cells expressing a CD4-targeting receptor selectively transferred cargo into CD4^+^ T cells (**Fig. 1l**). Similarly, donor cells expressing a FITC-targeting receptor and treated with FITC-conjugated anti-CD8 antibody selectively transferred cargo into CD8^+^ T cells (**Fig. 1m**). These findings demonstrate that trogocytosis-mediated transfer is programmable and specific across different cell types and endogenous ligands.

### Systematic characterization of receptor designs on molecular transfer outcomes

We investigated how the transmembrane domain of the receptor influence trogocytosis efficiency. We replaced the CD8 transmembrane domain with those from mouse Notch, human epidermal growth factor receptor (EGFR), and a glycosylphosphatidylinositol (GPI) membrane anchor (**Fig. 2a**). All transmembrane domains enabled efficient transfer (**Fig. 2b**), but the GPI anchor showed the highest transfer efficiency in terms of the amount of cargo per recipient cell (**Fig. 2c**).

**Figure 2:**
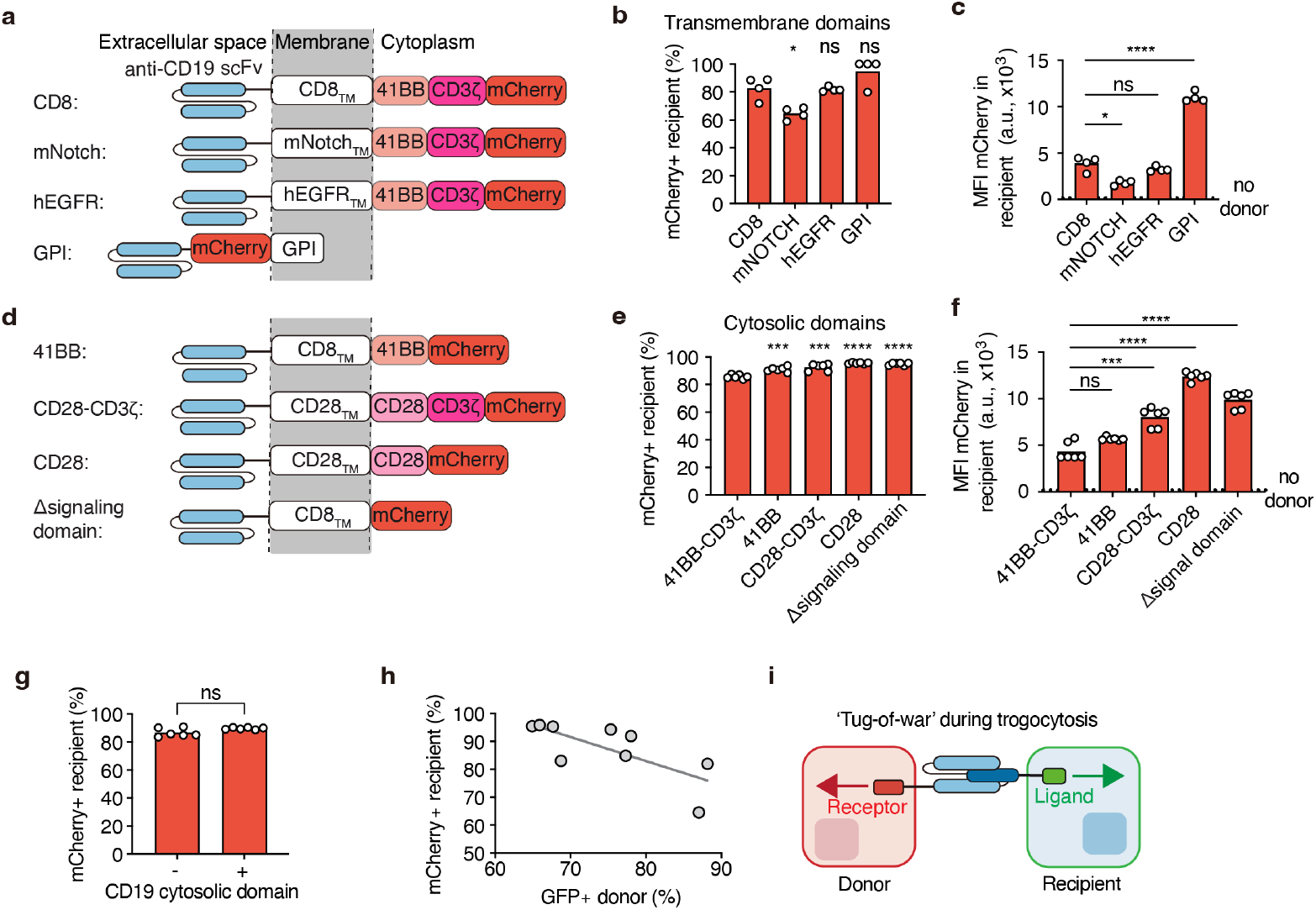
Characterization of receptor designs on molecular transfer outcomes. (**a**) Schematic showing modifications to the transmembrane domain (TM) of the receptor (originally CD8TM). (**b, c**) Percentage of mCherry^+^ and MFI mCherry in recipient cells after 24-hour 1:1 co-culture between Jurkat donor cells and CD19^+^ K562 recipient cells. Statistical tests were performed against the CD8TM group. n = 4 biological replicates. (**d**) Schematic showing modifications to the cytosolic domain of the receptor (originally 41BB-CD3ζ). (**e, f**) Percentage of mCherry^+^ and MFI mCherry in recipient cells after 24-hour 1:1 co-culture between Jurkat donor cells and CD19^+^ K562 recipient cells. Statistical tests were performed against the 41BB-CD3ζ group. n = 4 biological replicates. (**g**) Percentage of mCherry^+^ recipient cells in 24-hour 1:1 co-culture of anti-CD19 CAR-mCherry Jurkat and CD19^+^ K562. n = 6 biological replicates. (**h**) Correlation of the percentage of GFP^+^ Jurkat and the percentage of mCherry^+^ K562 upon 1:1 co-culture using different receptor architectures, based on mean values from 4 biological replicates. A simple linear regression was performed. R^2^ = 0.5361. (**i**) Schematic illustrating a ‘tug-of-war’ mechanism governing bidirectional transfer. Donor and recipient compete to acquire each other’s membrane fragment and associated molecules, resulting in an anti-correlation between the two directions. Welch’s two-sided t test. p<0.0001, ****; 0.0001≤p<0.001, ***; 0.001≤p<0.01, **; 0.01≤p<0.05, *; p≥0.05, non-significant (ns).

For the cytosolic domain, we tested 41BB, CD28, 41BB-CD3ζ, CD28-CD3ζ, and a truncated receptor lacking a signaling domain. All variants mediated efficient transfer (**Fig. 2d, e**). Among all architectures, receptors with the CD28 domain or no signaling domain showed the highest trogocytosis efficiency (**Fig. 2f**). Furthermore, we found that removing the signaling domain of the ligand CD19 in recipient cells, while preserving ligand expression levels, had no impact on trogocytosis (**Fig. 2g**), indicating that the signaling domain of the ligand is also dispensable.

Receptor architectures that reduced receptor transfer to recipient cells (*e*.*g*., mNOTCH) enhanced ligand transfer into donor cells, whereas architectures that promoted receptor transfer (e.g., Δsignaling domain and CD28) diminished the ability of donor cells to acquire opposing surface ligands (**Fig. 2h**). These findings suggest a “tug-of-war” mechanism during bi-directional trogocytosis (**Fig. 2i**).

### Cell surface fusogen enables functionalization of trogocytosed cargos

Although engineered receptors mediated ligand-specific protein transfer, microscopic images revealed that trogocytosed molecules colocalized with lysosomes in recipient cells (**Fig. 3a)**. To examine whether trogocytosed molecules are translocated to the cytoplasm for proper functionality, we developed a reporter recipient cell line stably expressing GFP11, a split GFP fragment that reconstitutes GFP upon binding with GFP1-10 (**Fig. 3b, c**). Although donor cells expressing anti-CD19 CAR fused to GFP1-10 successful transferred GFP1-10 through trogocytosis, no GFP signal was detected in recipient cells (**Fig. 3d**). These results indicate the need for further engineering to functionalize the trogocytosed molecules in recipient cells.

**Figure 3:**
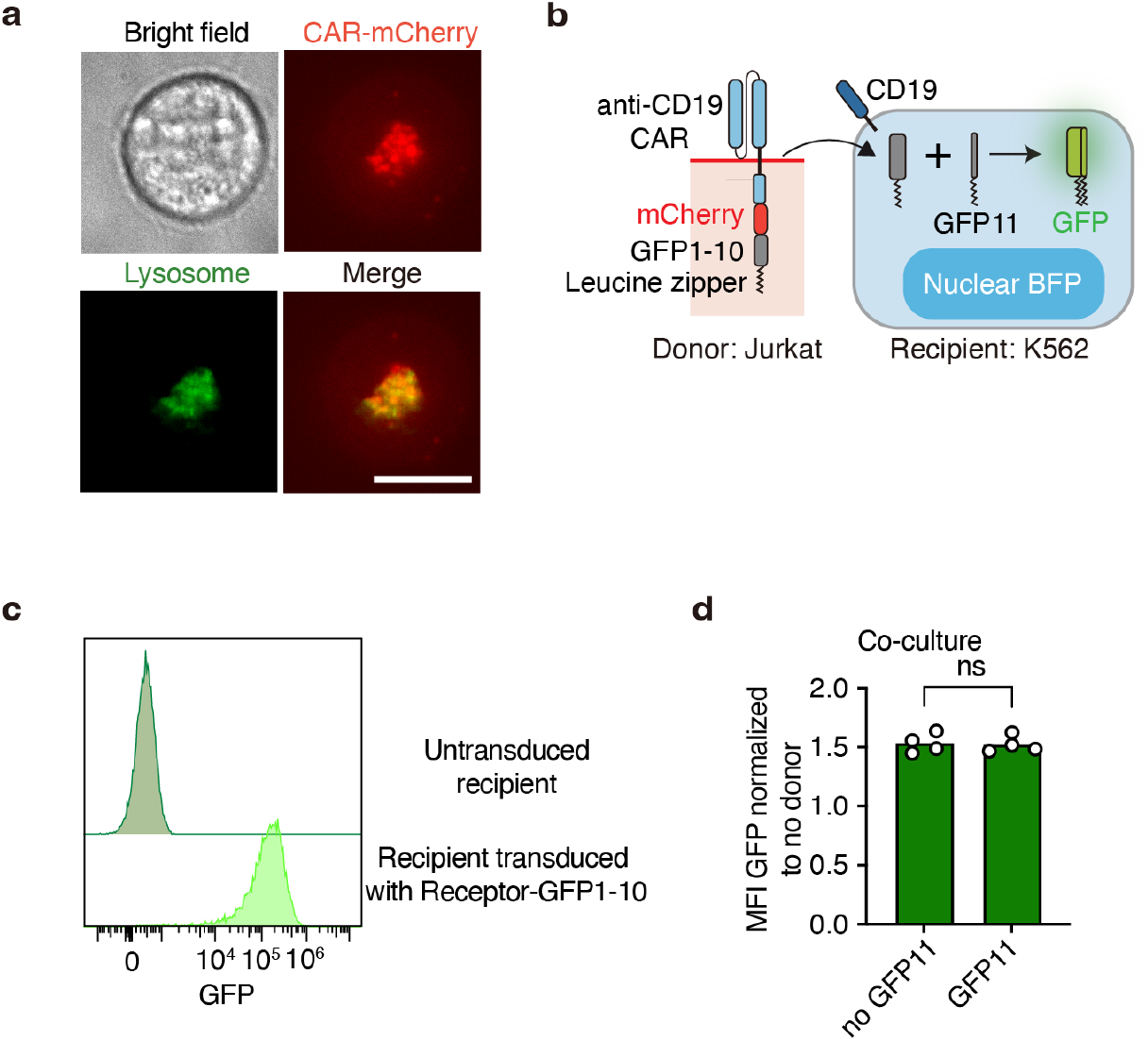
Transferred cargos undergo degradation without functionality upon trogocytosis. (**a**) Microscopy images of a CD19^+^ recipient cell co-cultured with anti-CD19 CAR-mCherry donor cell at a 1:1 ratio for 2-3 hours. Lysosomes were labeled with LysoView 488. Images show channels of the bright field, FITC (GFP), mCherry, and merged FITC-mCherry views. Scale bar, 10 μm. (**b**) Upon transfer into K562 recipient cells, if the receptor cytoplasmic domain GFP1-10 is exposed to the cytoplasm, it enables reconstitution of GFP on the cell membrane with GFP11 expressed in recipient cells. (**c**) GFP reconstitution in recipient cells expressing GFP11-NES after transduction of anti-CD19 CAR-mCherry-GFP1-10-NES. (**d**) Absence of GFP signal in recipient cells upon co-culture. Control recipient cells, which expressed CD19 but not GFP11, were used for comparison. n = 4 biological replicates. Welch’s two-sided t test. p<0.0001, ****; 0.0001≤p<0.001, ***; 0.001≤p<0.01, **; 0.01≤p<0.05, *; p≥0.05, non-significant (ns).

Given that trogocytosis involves the transfer of membrane fragments and associated molecules, we explored whether co-expressing low pH-dependent fusogens, glycoproteins that facilitate membrane fusion, on donor cells could functionalize trogocytosed molecules. We hypothesize that, upon internalization into the endo-lysosomal system, where the pH drops, co-transferred fusogens could promote fusion between trogocytosed membrane fragments and the endosome membrane, thereby releasing endosome-trapped molecules into the cytoplasm of recipient cells.

We co-expressed vesicular stomatitis virus glycoprotein (VSVg), a viral fusogen with low pH-dependent membrane fusion activity (*27, 28*), in donor HEK293T cells expressing anti-CD19 CAR-mCherry-GFP1-10, co-cultured with recipient K562 cells expressing cytoplasm-localized GFP11 (**Fig. 4a**). Co-delivery of VSVg enabled effective GFP reconstitution (**Fig. 4b**). The GFP reconstitution was cell contact-dependent and was blocked in transwell assays (**Fig. 4c**).

**Figure 4:**
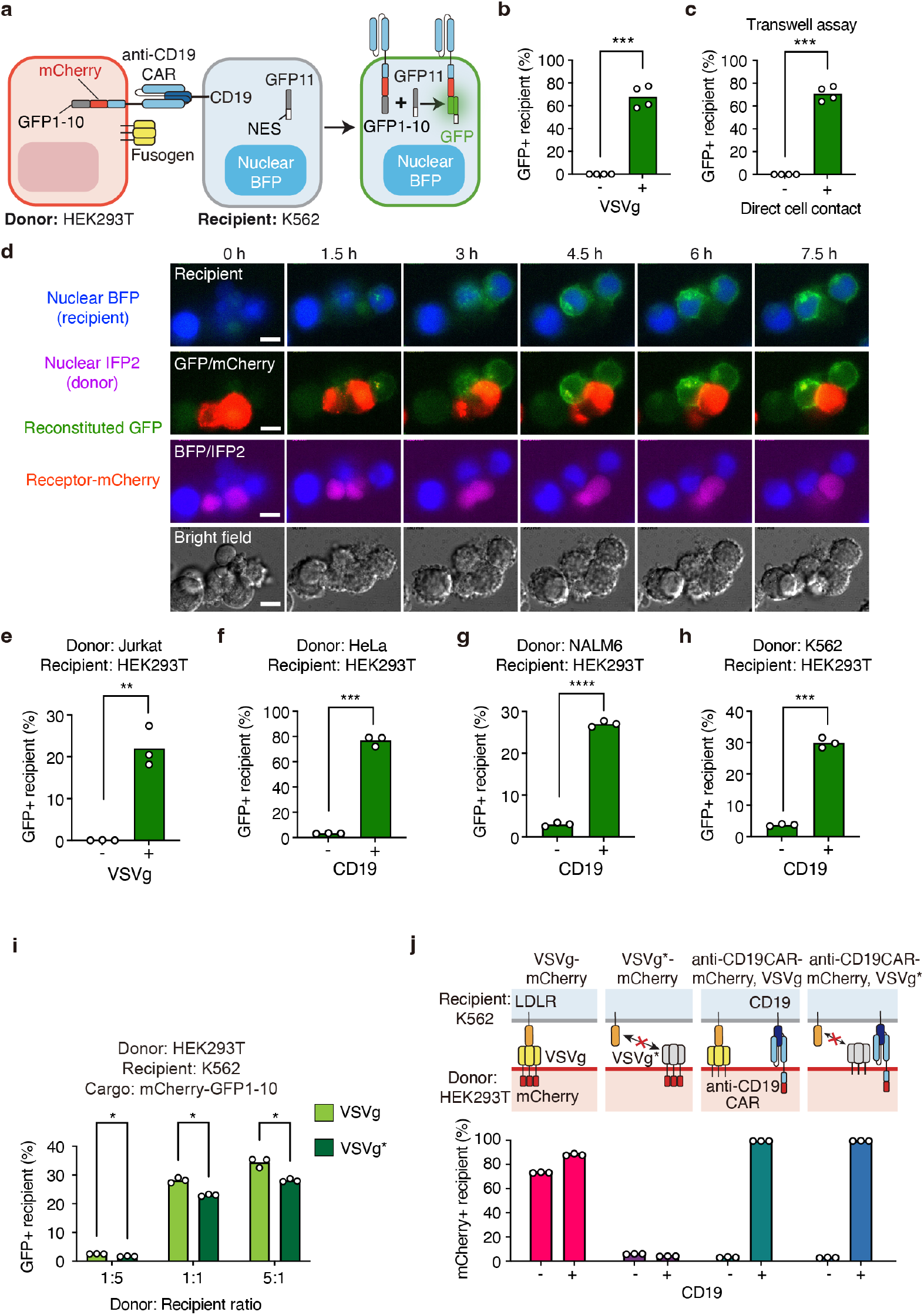
pH sensitive fusogen enables functionalization of transferred proteins in recipient cells. (**a**) Schematic illustrating how fusogens facilitate functional restoration of trogocytosed anti-CD19 CAR and mCherry-GFP1-10 cargo. Functionalized GFP1-10 associates with GFP11 in CD19^+^ K562 reporter cells to reconstitute GFP. NES, nuclear export signal. (**b**) Percentage of GFP^+^ recipient K562 cells among all recipient cells (BFP^+^) after 24-hour 5:1 co-culture with HEK293T donor cells stably expressing anti-CD19 CAR-mCherry-GFP1-10 and transfected with VSVg. The control group lacks VSVg transfection. n = 4 biological replicates across 2 independent experiments. (**c**) Transwell assay showing inhibition of cell-to-cell transfer using a 3 μm pore membrane separating donor and recipient cells. Positive group data overlaps with **b**, as they originate from the same experiment set. n = 4 biological replicates. (**d**) Microscopy images of VSVg-transfected HEK293T donor cells stably expressing anti-CD19 CAR-mCherry-GFP1-10 and nuclear IFP2, co-cultured 1:1 with CD19^+^ HEK293T recipient cells stably expressing GFP11-NES and nuclear BFP. Images show merged BFP/GFP (recipient), mCherry/GFP, BFP/IFP2, and the bright field. 0-hour marks the start of co-culture. Scale bar, 10 μm. (**e-h**) VSVg- and CD19-dependent functional receptor transfer from different donor cells to recipient cells. n = 3 biological replicates. (**i**) Donor HEK293T cells expressing anti-CD19 CAR-mCherry-GFP1-10 were transduced with either VSVg or VSVg* at equivalent expression levels. Donor cells were co-cultured with CD19^+^ K562 recipient cells expressing GFP11 for 24 hours, demonstrating comparable functional performance between VSVg and VSVg*. (**j**) Top, schematic of designs where mCherry were fused to VSVg, VSVg*, or anti-CD19 CAR co-expressed with VSVg or VSVg*. Percentage of mCherry^+^ recipient cells with 24-hour 1:1 co-culture with K562 cells. n = 3 biological replicates. Welch’s two-sided t test. p<0.0001, ****; 0.0001≤p<0.001, ***; 0.001≤p<0.01, **; 0.01≤p<0.05, *; p≥0.05, non-significant (ns).

Time-lapse microscopy revealed rapid mCherry accumulation in recipient cells within minutes of contact, followed by endosomal escape, which exposed GFP1-10 to the cytoplasm and enabled GFP reconstitution on the recipient plasma membrane within 2 hours (**Fig. 4d**). These findings highlight endosome membrane fusion and endosomal escape as critical steps in functionalizing trogocytosed molecules in recipient cells, without direct cell-cell fusion. These findings highlight endosome membrane fusion and endosomal escape as critical steps in functionalizing trogocytosed molecules in recipient cells. Similarly, multiple donor cell types engineered with a cognate receptor and VSVg successfully transferred functional GFP1-10 to targeted recipient cells (**Fig. 4e-h**).

To elucidate the role of the fusogen, we compared wildtype VSVg to the VSVg* variant (K47Q, R354A), which is deficient in low-density lipoprotein receptor (LDLR) binding (*29*). At similar expression levels, VSVg* exhibited comparable fusogenic activity to wildtype VSVg in mediating endosomal escape of trogocytosed molecules, indicating that LDLR binding is not a driver in functional trogocytosis (**Fig. 4i**). When mCherry was fused to the fusogen, wildtype VSVg-mCherry transferred into recipient cells independently of CD19, while VSVg*-mCherry did not, suggesting that LDLR binding by wildtype VSVg may facilitate off-target transfer of fusogen-coupled molecules (**Fig. 4j**). Despite that, neither VSVg nor VSVg* mediated non-specific transfer of anti-CD19 CAR-mCherry to CD19^-^ cells. To avoid non-specific VSVg-mediated transfer, we incorporated VSVg* into the final design.

### Cell-to-cell transfer enables delivery of non-membrane cargos including Cas9

Natural trogocytosis is primarily restricted to surface-bound molecules. Releasing trogocytosed molecules from the membrane could unlock ways to deliver proteins with diverse functions, including gene editing molecules, via cell-cell contact. We explored placing pH intein, a pH-sensitive self-cleavage domain (*30, 31*), between the ligand-binding domain and mCherry-GFP1-10 fused to a nuclear localization signal (NLS). To test the nuclear translocation of cargo, we adopted a reporter HEK293T cell line expressing nuclear GFP11 (**Fig. 5a**). With preferential self-cleavage of pH intein in the low pH environment of endosome, the cargo could translocate to the nucleus of recipient cells and reconstitute GFP. Indeed, GFP reconstitution was observed in the nuclei of recipient cells upon the addition of pH intein (**Fig. 5b**).

**Figure 5:**
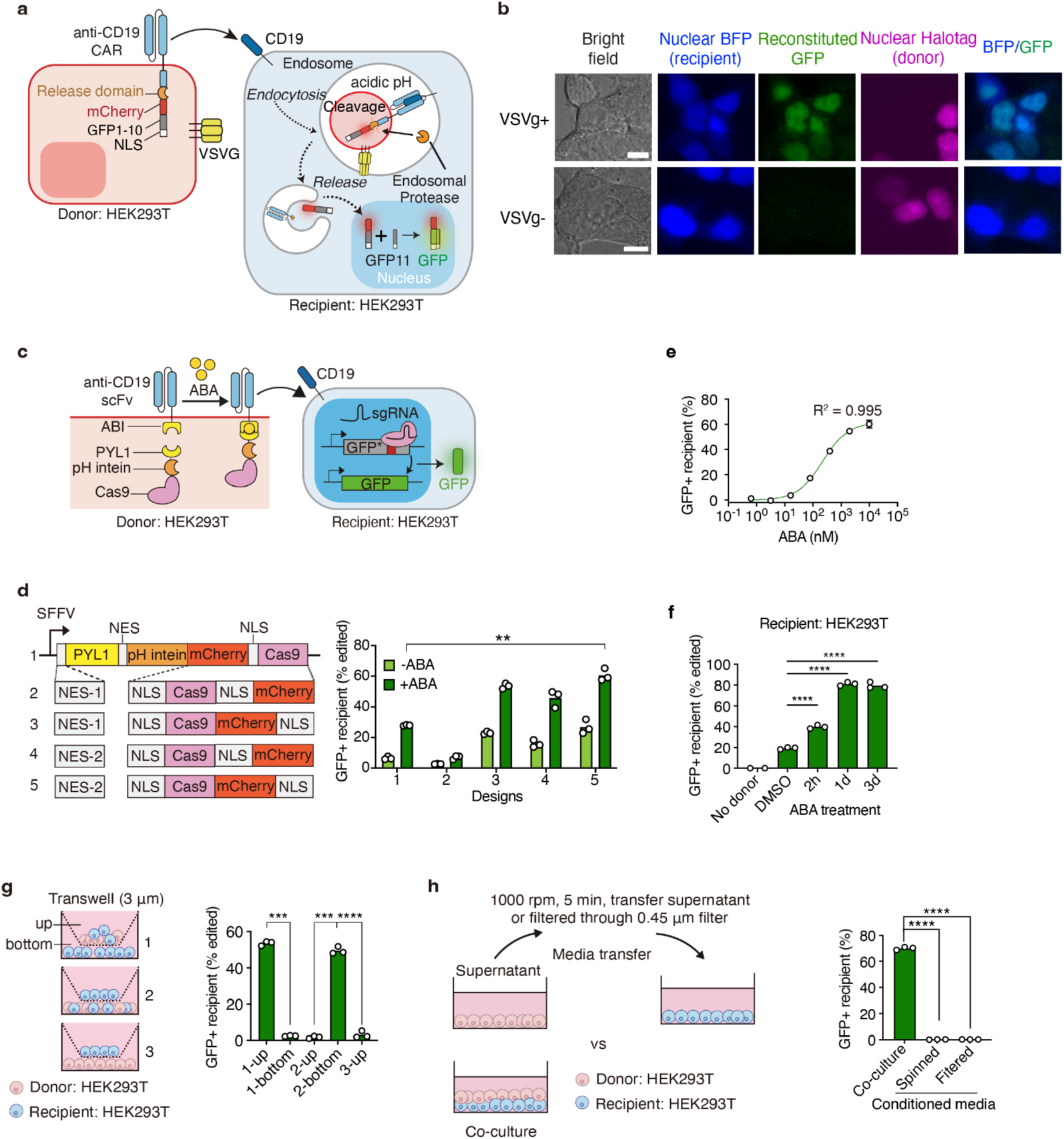
Investigation of molecular designs enabling translocation of transferred proteins to different compartments. (**a**) Schematic of constructs transduced into donor and recipient cells. A release domain was inserted between anti-CD19 CAR and mCherry-GFP1-10 with nuclear localization signal (NLS) fused to the C terminus. In recipient cells, NLS was also added to GFP11. Successful endosome escape, cleavage from membrane, and nuclear translocation lead to GFP reconstitution in the nucleus of recipient cells. (**b**) Microscopic images of VSVg-transfected anti-CD19 CAR-pH intein-mCherry-GFP1-10-NLS HEK293T donor cells and CD19^+^ GFP11-NLS HEK293T recipient cells with a 1:1 co-culture for 24 hours. The control group has no VSVg expression. Scale bar, 10 μm. (**c**) Schematic of an ABA-inducible Cas9 membrane localization system using ABI-PYL1 heterodimerization to anti-CD19 receptor in donor cells. Recipient cells encode a genomically integrated frameshifted GFP, an sgRNA targeting the reporter, CD19, and nuclear BFP. Functionalization of transferred Cas9 and subsequent nuclear translocation enable sgRNA-directed gene editing, restoring GFP expression. (**d**) Donor cells transfected with VSVg were co-cultured with CD19^+^ HEK293T recipient cells expressing a fluorescent reporter and sgRNA at a 10:1 ratio for 2 days with ABA (10 μM). Recipient cells were identified by nuclear BFP. Comparison of PYL1-pH intein-Cas9 constructs with different NLS and NES arrangements in donor cells. n = 3 biological replicates. (**e**) Percentage of GFP^+^ recipient cells at different ABA concentrations, calculated by subtracting values from the DMSO-treated group. VSVg-transfected HEK293T donor cells and recipient HEK293T cells shown in **c** are co-cultured at a 5:1 ratio for 72 hours. R^2^ indicates the goodness of fit to a Hill equation. n = 3 biological replicates. Error bars indicate mean ± SD. (**f**) Donor HEK293T cells were co-cultured with CD19^+^ HEK293T recipient cells containing the Cas9 cutting reporter at a 5:1 ratio for 3 days. ABA was added to the co-culture system for the indicated durations, followed by washout. n = 3 biological replicates. (**g**) Schematic of the transwell co-culture setup with a 3 μm pore size. VSVg^+^ donor cells transduced with anti-CD19-ABI and PYL1-pH intein-Cas9 were co-cultured with recipient cells containing a Cas9 reporter cassette, sgRNA, and nuclear BFP at a 5:1 donor-to-recipient ratio for 3 days. Cas9 transfer was observed only in cells in direct contact, while cells segregated by the transwell membrane did not exhibit Cas9 transfer. n = 3 biological replicates. (**h**) Donor HEK293T cells (VSVg*^+^) stably expressing nuclear mCherry, anti-CD19-ABI, and PYL1-pH intein-Cas9 are co-cultured 1:1 with recipient HEK293T cell expressing a GFP reporter and nuclear BFP for 72 hours (10 μM ABA). Recipient cells are gated as BFP^+^ mCherry^-^. Control groups use donor cell supernatant collected daily, spinned or filtered, and freshly added to recipient cells for 3 days. n = 3 biological replicates. Each dot represents one replicate culture well.

We explored Cas9 transfer into the recipient nucleus using the pH intein. Anti-CD19 CAR lacking cytosolic signaling domains (anti-CD19) was selected to target recipient cells. We utilized an abscisic acid (ABA)-induced heterodimerization system between ABI and PYL1 to recruit Cas9 to the plasma membrane (**Fig. 5c**). Recipient CD19^+^ HEK293T cells were engineered with an inactive GFP reporter and a targeting single guide RNA (sgRNA), where membrane-released Cas9-induced indels could restore GFP expression via a frameshift.

Through the optimization of nuclear export signals (NESs) and NLSs to balance nuclear export in donor cells and nuclear import in recipient cells, the ABA-inducible localization of Cas9 mediated significant gene editing in recipient cells (**Fig. 5d**). The Cas9 transfer system exhibited tunability with varying ABA concentrations (**Fig. 5e**) and treatment durations (**Fig. 5f**), highlighting the critical role of membrane protein localization in efficient trogocytosis-mediated transfer.

Importantly, recipient cells segregated from donor cells in transwell assays or sharing media without direct contact showed no GFP signal (**Fig. 5g, h**). These findings confirm the strict cell-contact dependency of the transfer and rule out mediation by gesicles or other diffusive vesicles (*28*). Building on our findings of contact-dependent, ligand-specific functional protein trogocytosis, we developed and named the system “<u>tr</u>ogocytosis-based tr<u>ans</u>fer and <u>f</u>unctional <u>e</u>ffector <u>r</u>elease” (TRANSFER). This system combines a minimized receptor, inducibly-membrane localized cargo, release domain, and fusogen mutant, establishing a versatile and novel modality for targeted molecular delivery.

### TRANSFER is a versatile and programmable protein delivery platform

A key limitation of current delivery modalities is their inability to recognize multiple ligands to guide delivery decisions (*12, 32, 33*). In contrast, TRANSFER enables simultaneous recognition of multiple surface ligands as inputs and delivers functional payloads as outputs, offering a programmable and adaptable delivery system.

On the input side, to demonstrate selective targeting of CD19 and CD3 double-positive cells, donor cells were engineered with an anti-CD19 synNotch receptor (*34*), which undergoes intramembrane proteolysis upon CD19 recognition, releasing a nucleus-localized transcription factor. The transcription factor drives expression of a cargo fused to PYL1, which is subsequently loaded onto an anti-CD3 TRANSFER receptor via ABA-induced dimerization. This design effectively implemented an AND logic gate, enabling selective cargo delivery to target cells by sensing two inputs (**Fig. 6a, b**). Such programmable logic for multi-ligand sensing surpasses the capacities of current delivery vehicles.

**Figure 6:**
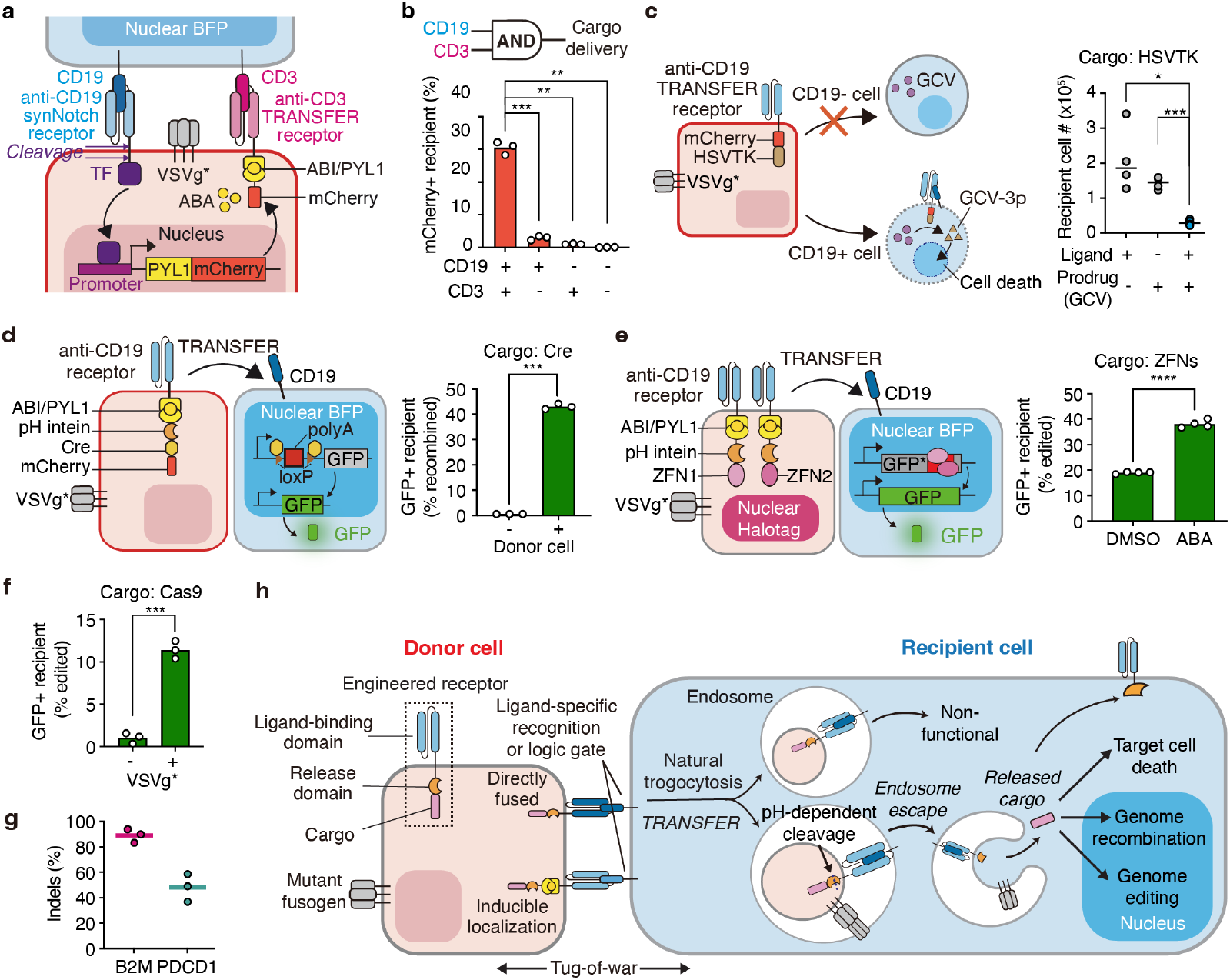
TRANSFER is a programmable delivery platform enabling combinatorial marker sensing. (**a-b**) TRANSFER enables ligand-specific delivery guided by an AND gate logic. Donor HEK293T cells expressing an anti-CD19 synNotch receptor fused to a transcription activator (GAL4-VP64), PYL1-mCherry driven by GAL4 binding sites (GAL4-UAS), anti-CD3-ABI, and VSVg* were co-cultured with four types of recipient cells (CD19^+^/CD3^+^ Jurkat, CD19^-^/CD3^+^ Jurkat, CD19^+^/CD3^-^ K562, CD19^-^/CD3^-^ K562) at a 1:1 ratio for 24 hours. Percentages of mCherry^+^ recipient cells are shown for each co-culture. Recipient cells were identified by nuclear BFP. n = 3 biological replicates. (**c**) Ligand-specific delivery of enzymes for targeted cell death. Left, schematic of donor HEK293T cells expressing anti-CD19-mCherry-HSVTK and VSVg*, co-cultured 5:1 with CD19^+/-^ recipient K562 cells for 3 days. Right, quantification of recipient cell numbers under different ligand and prodrug (GCV) conditions. GCV was added 1 day after co-culture initiation. Recipient cells were identified by nuclear BFP. n = 4 biological replicates. (**d**) Delivery of Cre recombinase for gene editing. Left, schematic of donor HEK293T cells expressing anti-CD19-ABI and PYL1-pH intein-Cre-mCherry-DHFR and VSVg*, co-cultured 1:1 with CD19^+^ recipient K562 cells expressing loxP-polyA-loxP-GFP for 2 days. Cre is fused to DHFR to mitigate toxicity in donor cells, with expression stabilized by adding 100μM trimethoprim (TMP). Right, percentages of GFP^+^ recipient cells with and without co-culture. n = 3 biological replicates. (**e**) Delivery of ZFNs for genome editing. Left, donor HEK293T cells were transduced with anti-CD19-ABI and nuclear HaloTag and transfected with PYL1-pH intein-ZFN1, PYL1-pH intein-ZFN2, and VSVg*. Recipient cells were transduced with a frameshifted GFP reporter with ZFN target sites, enabling ZFN-mediated GFP restoration. Right, percentages of GFP^+^ recipient cells after co-culture 5:1 with donor cells for 3 days (ABA 1 mM). Recipient cells were identified as BFP^+^ Halotag^-^. n = 4 biological replicates. (**f**) Transferred Cas9-mediated reporter editing. Percentage of GFP^+^ recipient Jurkat cells co-cultured 10:1 with donor HEK293T cell expressing anti-CD3-ABI, PYL1-pH intein-Cas9, nuclear mCherry, and VSVg* for 3 days (ABA 1mM). Recipient cells expressed nuclear BFP, an inactive GFP reporter with a sgRNA-targeting site, and sgRNA. n = 3 biological replicates. (**g**) Trogocytosed Cas9-mediated endogenous gene editing. Indel percentages for Cas9-mediated editing at B2M or PDCD1 loci. Donor HEK293T cells expressing anti-CD19-ABI, PYL1-pH intein-Cas9, nuclear mCherry, and VSVg*, were co-cultured 1:1 with CD19^+^ recipient HEK293T cells expressing nuclear BFP and sgRNA targeting B2M or PDCD1 for 3 days (ABA 10 μM). Amplicon sequencing was performed on sorted BFP^+^ mCherry^-^ recipient cells. n = 3 biological replicates. (**h**) Summary of the TRANSFER mechanism as a versatile, programmable, and specific delivery system applicable to various cell types and functional cargos. Welch’s two-sided t test. p<0.0001, ****; 0.0001≤p<0.001, ***; 0.001≤p<0.01, **; 0.01≤p<0.05, *; p≥0.05, non-significant (ns).

On the output side, TRANSFER can be applied to deliver cytoplasmic functional cargo, such as Herpes simplex virus thymidine kinase (HSVTK), a prodrug-converting enzyme that phosphorylates the non-toxic drug Ganciclovir (GCV) into GCV-triphosphate (GCV-3p), inducing cell death in recipient cells (*35*). Fusing HSVTK to the anti-CD19 receptor in donor cells enabled specific transfer to CD19^+^ recipient cells, triggering targeted cell death, while CD19^-^ cells remained unaffected (**Fig. 6c**).

For cargos functioning in the nucleus, we found that Cre recombinase transfer effectively mediated recombination at loxP sites, generating GFP signal in reporter K562 cells (**Fig. 6d**). To confirm the ability of TRANSFER to deliver multiple cargos simultaneously, we tested a pair of zinc finger nucleases (ZFNs) (*36*). Donor HEK293T cells engineered with the ZFN pair efficiently transferred them to recipient HEK293T cells encoding genomically integrated, frameshifted GFP, restoring GFP expression via ZFN-mediated genome editing in an ABA-dependent manner (**Fig. 6e**).

TRANSFER-mediated Cas9 transfer could be generalized to other recipient cell types, including Jurkat, by targeting its endogenous ligand CD3 (**Fig. 6f**). In co-culture systems, trogocytosed Cas9 effectively edited endogenous genes, such as B2M and PDCD1 (**Fig. 6g**). These results establish the versatility of TRANSFER as a functional protein delivery platform.

## Discussion

As a natural cell-cell communication pathway, trogocytosis allows immune cells to acquire functional proteins they do not endogenously express from other cells (*37*–*39*). Building on this natural process, we investigated engineered trogocytosis as a versatile, specific, and cell contact-dependent molecular transfer mechanism (**Fig. 6h**).

Compared to non-living delivery vehicles, engineered cells offer unique advantages, including spatiotemporal control of cargo production *in situ*, sensing and responding to multiple inputs (*34*), and potential active migration across biological barriers (*40*). Achieving specific and programmable delivery between living cells is essential to realizing this potential. Previous approaches, such as engineering cells to produce designer exosomes (*41*) or RNA exporters (*42*), have primarily focused on mRNA delivery and face limitations such as poor cell type specificity (*42*) or rapid clearance by the spleen and liver (*41*). In contrast, TRANSFER leverages a contact-based mechanism to achieve specific and programmable protein delivery, avoiding the off-target uptake associated with diffusive secreted vesicles.

In addition to trogocytosis, other mechanisms of trans-synaptic material transfer have been documented, including trans-synaptic vesicles (*43, 44*), which are extracellular vesicles formed by T cells at the immunological synapse to transport molecules across the synaptic cleft(*45*), and trans-endocytosis, the endocytosis of cellular materials by another cell (*46*). As there is no specific and universal pharmacological inhibitor of trogocytosis (*7, 19*), and donor cell membrane budding could potentially cooperate with trogocytosis in cell-to-cell transfer (*38, 45, 47*), we cannot rule out the possibility that the trans-synaptic transfer observed here involves a combination of trogocytosis, trans-endocytosis, and trans-synaptic vesicles, with possible variations in the mechanism depending on the cell types used.

Overall, our work establishes a novel and universal cell-to-cell transfer mechanism, uncovering its unique principles and demonstrating its adaptability across diverse cell types and molecular cargos (**Fig. 6h**). The TRANSFER platform provides a foundation for programmable and versatile cell-based functional molecular delivery in targeted therapeutics and biotechnological applications.

## Acknowledgments

The authors thank all members of the Lei Stanley Qi lab for facilitating experiments and helpful discussions, Xiaoshu Xu for sharing the Cas9 reporter construct, Mengting Han and Yanyu Zhu for assistance with imaging, Yitong Ma, Jon Bezney, Victor Tieu and Michael Chavez for helpful discussions on experiments and computational analysis. We thank Aditi Merchant for technical support. The authors thank Crystal Mackall and Robbie Majzner (Stanford University) for sharing the CAR constructs, Jiangbin Ye and Bo He (Stanford University) for providing the MCF7 cell line, Liqun Zhou (Yale University) for sharing the tumor digestion protocol. We thank Wendell Lim (UCSF), Rogelio Hernández-López and Jeremy Villafuerte (Stanford University) for providing the SynNotch plasmids, and Sui Wang, Bo Xiong, Nirk Ericson Quispe Calla and Cesar Walter Guzman Huancas (Stanford University) for their advice on animal work. We thank Takuji Iwasato (National Institute of Genetics), Michael Davidson (UCSD), and Xiaokun Shu (UCSF) for Addgene plasmids. The authors thank Alice Ting (Stanford University), Michael McManus (UCSF), and Daniel Fletcher (UC Berkeley) for helpful discussions.

X.C. acknowledges support by the Stanford Bio-X SIGF Fellowship. L.S.Q. acknowledges support by the Chau Hoi Shuen Foundation, National Science Foundation (NSF) CAREER award, National Institutes of Health (NIH), and California Institute for Regenerative Medicine (CIRM). This work is supported by Targeted Genome Editor Delivery (TARGETED) Challenge (L.S.Q), National Science Foundation CAREER award 2046650 (L.S.Q.), National Institutes of Health grant R01CA266470 and R21AG077193 (L.S.Q.), NIH Director’s Pioneer Award DP1NS137219 (L.S.Q.). L.S.Q. is a Chan Zuckerberg Biohub – San Francisco investigator.

